# Tapered Drug delivery, Optical stimulation, and Electrophysiology (T-DOpE) probes reveal the importance of cannabinoid signaling in hippocampal CA1 oscillations in behaving mice

**DOI:** 10.1101/2023.06.08.544251

**Authors:** Jongwoon Kim, Hengji Huang, Earl Gilbert, Kaiser Arndt, Daniel Fine English, Xiaoting Jia

## Abstract

Understanding the neural basis of behavior requires monitoring and manipulating combinations of physiological elements and their interactions in behaving animals. Here we developed a thermal tapering process (TTP) which enables the fabrication of novel, low-cost, flexible probes that combine ultrafine features of dense electrodes, optical waveguides, and microfluidic channels. Furthermore, we developed a semi-automated backend connection allowing scalable assembly of the probes. We demonstrate that our T-DOpE (**T**apered **D**rug delivery, **Op**tical stimulation, and **E**lectrophysiology) probe achieves in a single neuron-scale device (1) high-fidelity electrophysiological recording (2) focal drug delivery and (3) optical stimulation. With a tapered geometry, the device tip can be minimized (as small as 50 μm) to ensure minimal tissue damage while the backend is ~20 times larger allowing for direct integration with industrial-scale connectorization. Acute and chronic implantation of the probes in mouse hippocampus CA1 revealed canonical neuronal activity at the level of local field potentials and spiking. Taking advantage of the triple-functionality of the T-DOpE probe, we monitored local field potentials with simultaneous manipulation of endogenous type 1 cannabinoid receptors (CB1R; via microfluidic agonist delivery) and CA1 pyramidal cell membrane potential (optogenetic activation). Electro-pharmacological experiments revealed that focal infusion of CB1R agonist CP-55,940 in dorsal CA1 downregulated theta and sharp wave-ripple oscillations. Furthermore, using the full electro-pharmacological-optical feature set of the T-DOpE probe we found that CB1R activation reduces sharp wave-ripples (SPW-Rs) by impairing the innate SPW-R-generating ability of the CA1 circuit.

## Main

Recent advancements in neural interface technology have significantly improved our understanding of the nervous system,^1-10^ presenting new opportunities for the treatment of neurological disorders and the development of brain-machine interfaces. However, the complexity of the brain’s inner workings demands increased precision in the monitoring and manipulation of neural activity in order to decipher and understand its connectivity and dynamics.^1, 11-14^ Optogenetics has been developed as a powerful tool to modulate and/or monitor neural activities with a high spatiotemporal resolution and cell-type specificity.^15, 16^ It has led to the widespread adoption of bi-directional devices, which are able to monitor electrical activity and apply optical stimulation.^4, 17-20^ However, other biological factors such as neurochemistry are intertwined with electrical activity, necessitating simultaneous opto-electro-pharmacological investigations, which is hard to achieve with current devices. Therefore, developing single devices that can interact with the brain of behaving mammals across such multiple modalities is a critical goal for the field.

A few technologies have been developed aiming to address this challenge;^21, 22^ however, the fabrication of these probes is mostly based on cleanroom microfabrication techniques which are time-consuming and expensive. As an alternative, thermal fiber drawing is a promising technique for producing scalable multimodality fiber devices at a low cost.^23-29^ Such fiber devices are fabricated via a method commonly used in industry to produce optical fibers. The macroscale, multi-material preform is heated until softened, and pulled into hundreds of meters of fibers that can be as thin as a human hair. The fast and simple fabrication process utilizing affordable machinery and soft material results in a cheap, sturdy and biocompatible device. However, an inherent challenge of fiber technology lies in the uniform diameter across the length: there is a tradeoff between minimized sensing tip for biocompatibility and maximized backend tip for easy connection. This has limited the fiber’s practicality in neuroscience applications. To address this issue, we developed a novel thermal tapering process (TTP) and a semi-automated connection method, which enable us to fabricate microprobes with high structural and functional complexities and scalability (**fig. 1**, left). This novel **T**apered **D**rug delivery, **O**ptical Stimulation, and **E**lectrophysiology (T-DOpE) probe allows us to investigate highly complex neural circuitry such as the hippocampus of behaving mice.

**Figure 1:**
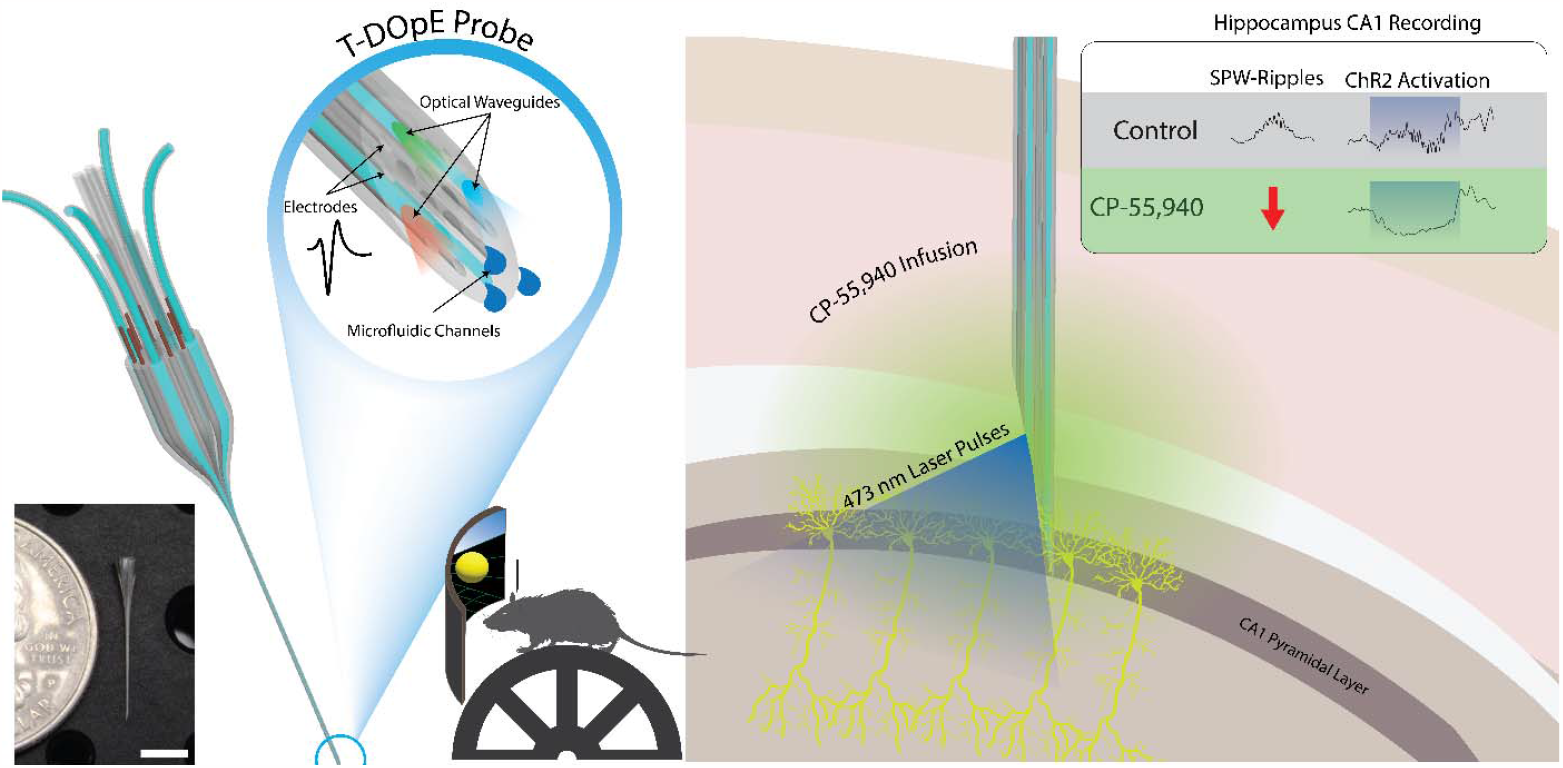
Illustration of the newly developed T-DOpE probe and CA1 electrophysiological activity in response to local chemical manipulations under natural and optically stimulated conditions. T-DOpE probe offers higher complexities at the sensing tip while easing the connection between the backend and the electronics. The probe is implanted into behaving head-fixed mice expressing channel rhodopsin (ChR2) in pyramidal neurons in hippocampal area CA1. Local infusion of synthetic cannabinoid (CB1 Agonist, CP-55,940) in CA1 is sufficient to abolish both spontaneous and optogenetically induced sharp wave-ripples (SPW-R) (Scale bar: 5mm).

Hippocampal circuit activity is critical for episodic and spatial memory. Hippocampal theta (~6-10 Hz), gamma (~35-80 Hz), and sharp wave-ripple (SPW-R, ~100-250 Hz) oscillations all contribute to mnemonic functions of the circuitry.^30-34^ In rodents, these oscillations are disrupted following systemic pharmacological activation of cannabinoid receptors by compounds such as Δ-9-tetrahydrocannabinol (Δ9-THC) or agonists of cannabinoid receptors (CB1Rs). This is suggested to be a mechanism behind cannabinoid-associated memory impairment in rodents^35-38^ and humans.^39-41^ It is believed that activation of CB1Rs impairs memory by changing the activity of hippocampal neurons expressing CB1Rs as well as their synaptic partners in local hippocampal circuits.^37, 42-45^ The synthetic cannabinoid, CP-55,940, is a useful tool to study the effect of CB1R activation in rodent models. Previously, it has been shown systemic administration of CP-55,940 in rats most prominently reduces theta oscillations and SPW-Rs.^37^ Importantly, systemic administration of cannabinoids cannot rule out CA1 interactions with other brain areas. The same study also found that in urethane-anesthetized rats, intrahippocampal delivery of CP-55,940 abolished SPW-Rs,^37^ suggesting that the effects of systemic administration may be mediated by intra-hippocampal changes in CB1R signaling. The effect of focal CA1 CB1R agonism has never been investigated in behaving animals, and because the only intrahippocampal administration was completed under anesthesia, the neuronal substrates and mechanisms by which CB1Rs control hippocampal rhythms remains unknown. Using our novel device, we investigated the role of CA1 CB1Rs in hippocampal local field potential activity using simultaneous optogenetic manipulation of CA1 neuron excitability and CB1R agonism via pharmacological intervention.

Here, we present the Thermal Tapering Process (TTP) which enables high feature density, functional complexity, and semi-automated backend connections. Various designs of the probe were fabricated to demonstrate the capabilities achievable with the tapering method. To test our device performance *in vivo*, T-DOpE probes were implanted in awake, behaving mice. We demonstrate that T-DOpE probes can reliably record hippocampal circuit electrophysiology in both acute and chronic platforms. We then demonstrated precisely controlled optical and chemical modulation using our probes in CA1. We confirmed and extended the understanding of cannabinoid effects in CA1 in awake and behaving mice. We found that intrahippocampal administration of CP-55,940 during head-fixed exploration was sufficient to reduce theta oscillations and abolish spontaneous SPW-R. Furthermore, we discovered that the local infusion of CP-55,940 in CA1 of a mouse expressing channel rhodopsin (ChR2) in pyramidal cells inhibits optogenetically induced SPW-Rs (**fig. 1**, right). For the first time, we used a single novel device, with unique multi-modal capabilities, to examine the effects of CA1 cannabinoid receptor activation on local oscillatory activity. Our findings reveal the importance of CB1Rs expressed by CA1 neurons in the generation of SPW-Rs.

## Result

### T-DOpE Probe fabricated using Thermal Tapering Process

Multifunctional tapered probes with various designs were fabricated via the thermal drawing process (TDP) followed by a novel thermal tapering process (TTP). **Fig. 2a** shows the fabrication steps for a preform. A Polycarbonate (PC) rod was machined to create hollow channels (microfluidic channels in the finished device), and spaces to inlay waveguides and electrodes. Waveguides with PC core (refractive index n =1.586) and Poly(methyl 2-methylpropenoate) (PMMA; refractive index n = 1.49) cladding and bismuth tin (BiSn) alloy electrodes were then inserted into their respective positions. The preform was then wrapped in PC film and consolidated in a vacuum furnace. Through TDP, the finalized preform was heated and drawn down to a 2mm diameter fiber (**fig. 2b**). The fiber diameter was closely monitored via a laser micrometer and controlled by adjusting the pulling speed and temperature. The fiber was cut into 10cm long mini-preforms (approximately hundred per TDP draw) for the subsequent thermal tapering process (TTP). During TTP, mini-preforms are heated, softened and pulled to form a tapered structure (**fig. 2c**). The pulled structure was then cut at a specific angle, resulting in two individual microprobes with 2mm diameter backends and 150μm tapered tips for monitoring and manipulating neural activity. For this study, three different probe designs (8/1/1, 8/4/8 and 0/8/12; electrodes/waveguides/microfluidic channels, respectively) were fabricated. Their device tip cross sections are shown in **fig. 2d**. The BiSn electrodes record extracellular voltage while the optical waveguides control optogenetics, and the microfluidic channels enable focal drug infusion.

**Figure 2:**
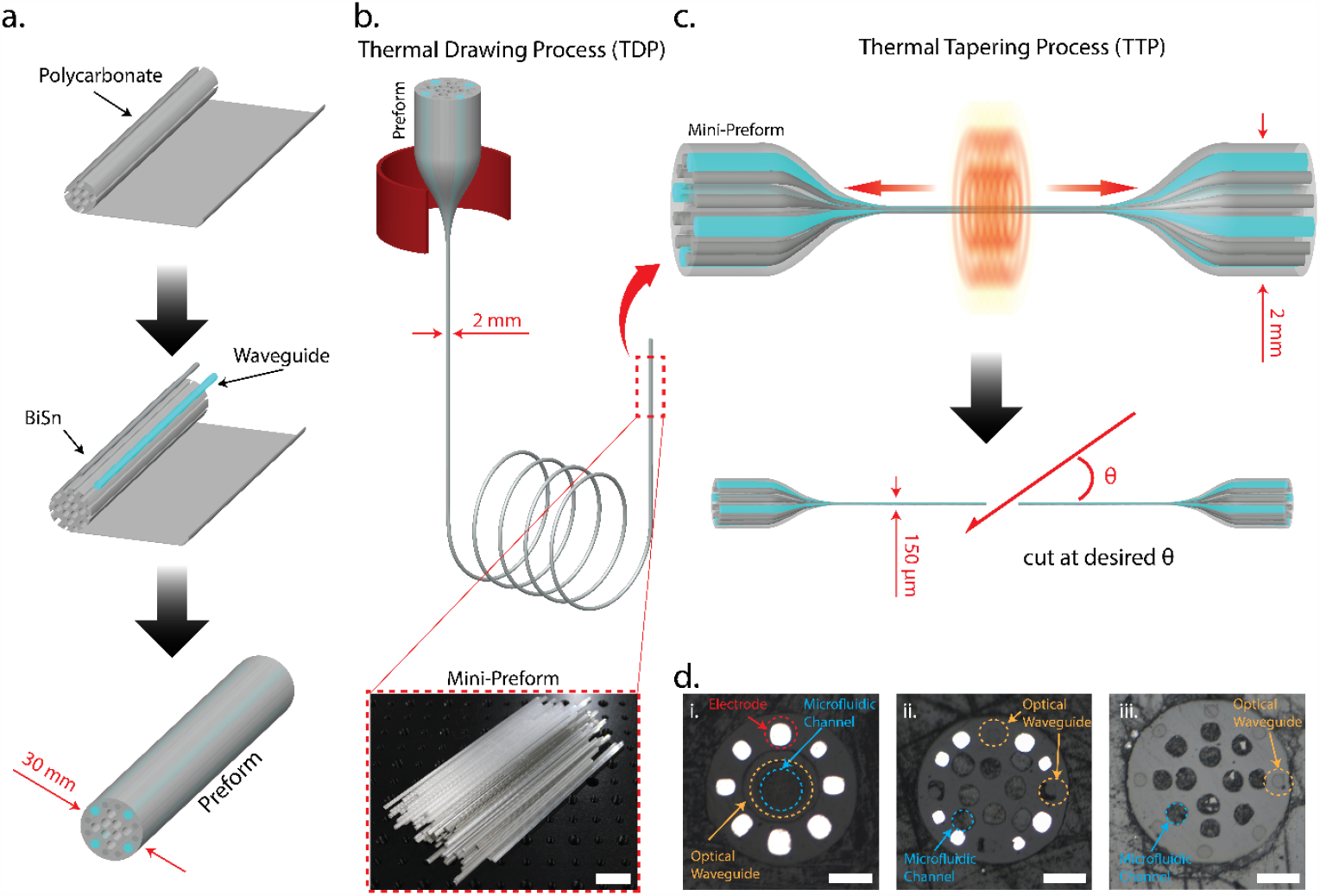
**a**. Schematic of the preform fabrication. **b**. Thermal drawing process pulling 30mm diameter preform into 2mm diameter fibers. The fibers are dissected into 10cm long mini-preforms for tapering (Scale bar: 2cm) **c**. Illustration of the thermal tapering process of mini preform. Similar to glass pipette pulling, the mini preform is heated until softened, pulled and cut at a desired angle, resulting in two individual probes. **d**. Cross-sectional images of the three various designs. **(i)** eight electrodes, one microfluidic channel, and one optical waveguide. **(ii)** Eight electrodes, eight microfluidic channels, and four optical waveguides. **(iii)** Twelve microfluidic channels and eight optical waveguides. (Scale bar: 50μm)

### T-DOpE probe connections and characterizations

We developed a novel fast, semi-automated backend connection method to replace the traditional method which is slow and labor-intensive. Our connection to the T-DOpE probe was executed exclusively on its 2mm backend (**fig. 3a**). Electrical connection was accomplished by heating the probe’s backend and inserting insulated copper wires into melted electrodes. The probe was then cooled for the BiSn electrodes to solidify. The microfluidic connection was achieved by inserting a custom drawn thin PC tube into the microfluidic channels. Custom drawn polymer optical fibers were coupled onto the waveguides on the probe’s backend. UV epoxy was then used to seal and fix the microfluidic and optical connections in place. A fully functional and connectorized probe with 8 electrodes, 4 waveguides and 8 microfluidic channels is shown in **fig. 3b**. Probes are fitted with industry standard adapters compatible with electrophysiology equipment, optical modules, and drug delivery pumps. A photograph of a probe connected to a printed circuit board (PCB) is shown in **Supplementary fig**.**1**.

**Figure 3:**
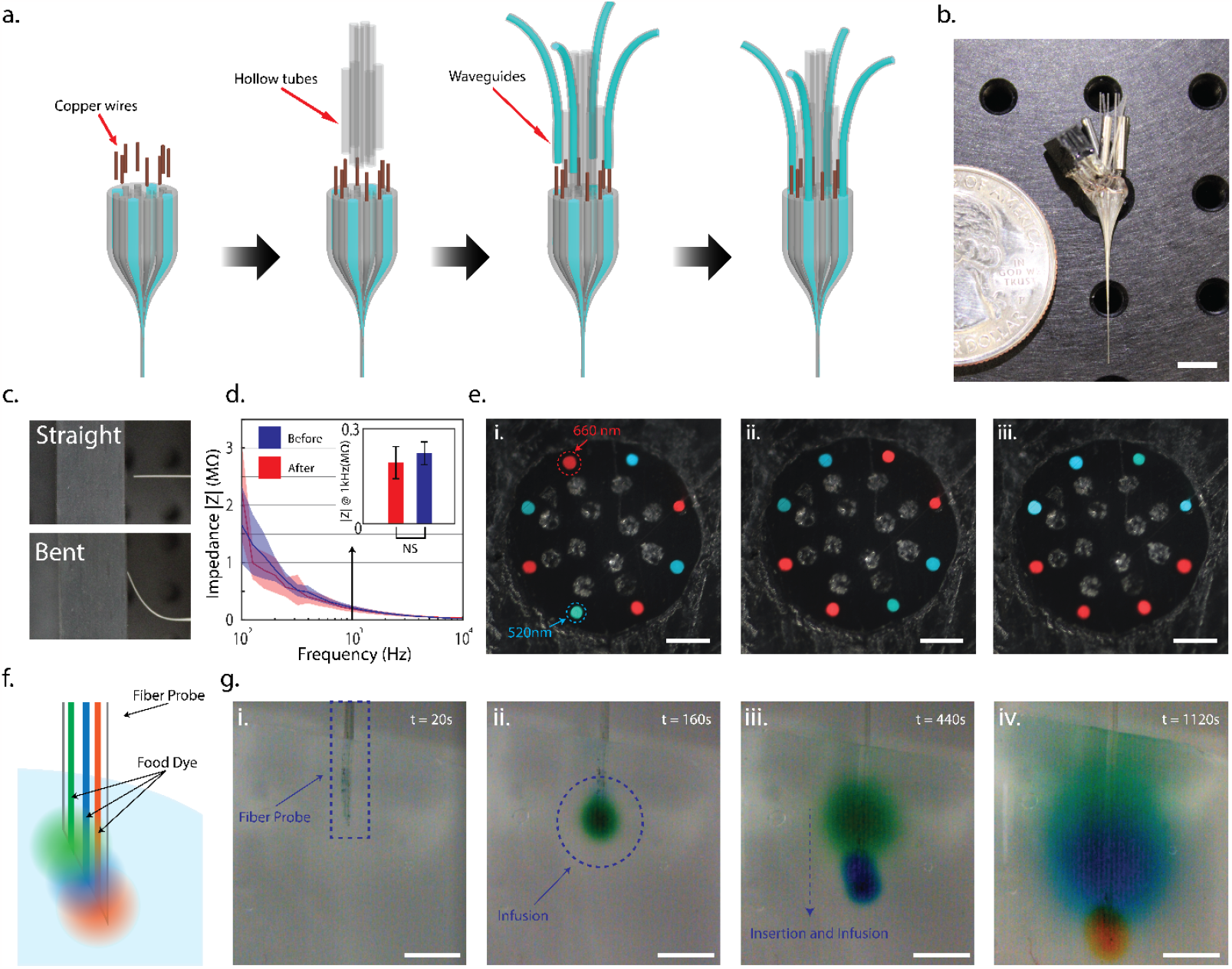
**a**. Schematic of the T-DOpE probe backend connection process. **b**. Photograph of a fully connected probe with eight electrodes, eight microfluidic channels, and four optical waveguides. (Scale bar: 5mm) **c**. Photograph of a straight probe and a probe bent roughly 45° to demonstrate the flexibility at the sensing end. **d**. Impedance measurements of the BiSn electrodes before and after bending. All error bars and shaded colors represent the standard deviation. (Student’s two-tailed t-test, NS: not significant, p>0.05(After vs. Before, p=0.3717, n=8)) **e**. Crossectional images to illustrate the individual addressability of the optical waveguides. (Scale bar: 50μm) **f**. Cartoon of green, blue, and orange food dye independently injected via the probe into a brain phantom. **g**. Time-lapsed images to demonstrate the drug infusion in a 0.6% agarose gel. The inserted probe demonstrates the infusion of three different dyes at three different heights in the phantom. (Scale bar: 1mm)

To demonstrate the flexibility and durability of the probe, we bent the probe at a >45° by applying axial force pressing the tip on a hard surface as shown in **fig. 3c**. We measured the spectral impedance of the T-DOpE probe before and after bending. As shown in fig. 3d, bending has no statistical impact on our device’s impedance at 1 kHz, with an average impedance of 192 ± 50 kΩ before bending and 220 ± 30 kΩ after bending. (Student’s two-tailed t-test, NS: not significant, p>0.05 (After vs. Before, p=0.3717, n=8)). We also demonstrated our optical waveguide’s individual addressability in our device with 8 waveguides and 12 microfluidic channels. The cross-sectional images of the device with red and green lights emitting from the waveguides are shown in **fig. 3e**. To test our device’s ability to deliver multiple drugs independently via separate microfluidic channels, we conducted an experiment with a device of the same design in 0.6% agarose gel as illustrated in **fig. 3f**. The device was first inserted in the agarose gel and delivered green food dye (**fig. 3g**). The blue and orange dye were infused after insertion into deeper regions of the gel. The diffusion of the dyes clearly indicated the infused regions of the agarose gel. These measurements and tests verify the functionality of our probe for extracellular recording, optogenetic manipulation, and drug delivery.

### *In vivo* capabilities of T-DOpE probe: electrophysiology recording, optogenetic, and drug infusion

To evaluate the functionalities of the T-DOpE probe *in vivo*, we collected acute and chronic electrophysiological activity in behaving mice. As shown in **fig. 4a**, probe implantation was targeted to the CA1 region of the hippocampus. **Fig. 4b** shows an example of an acute wideband (0.1 – 8000 Hz) extracellular trace from CA1, which demonstrates a transition from rest, when SPW-Rs occur, to a running state, when theta oscillations emerge. Shaded region **i**. demonstrates multi-unit activity in the raw trace (i.e., bandpass filtered at 0.1-8000 Hz). Shaded regions **ii**. and **iii**. indicate oscillations which are established as identifying electrophysiological landmarks of hippocampus CA1^30-32^: SPW-Rs (**ii**., 100-250 Hz), and theta-nested gamma oscillations (**iii**., Theta: 6-11 Hz, Gamma: 40-80 Hz)^33, 34^. Our new T-DOpE probe can thus acquire single units and local field potential (LFP) activity in behaving animals without disrupting natural physiological process within the local circuit.

**Figure 4:**
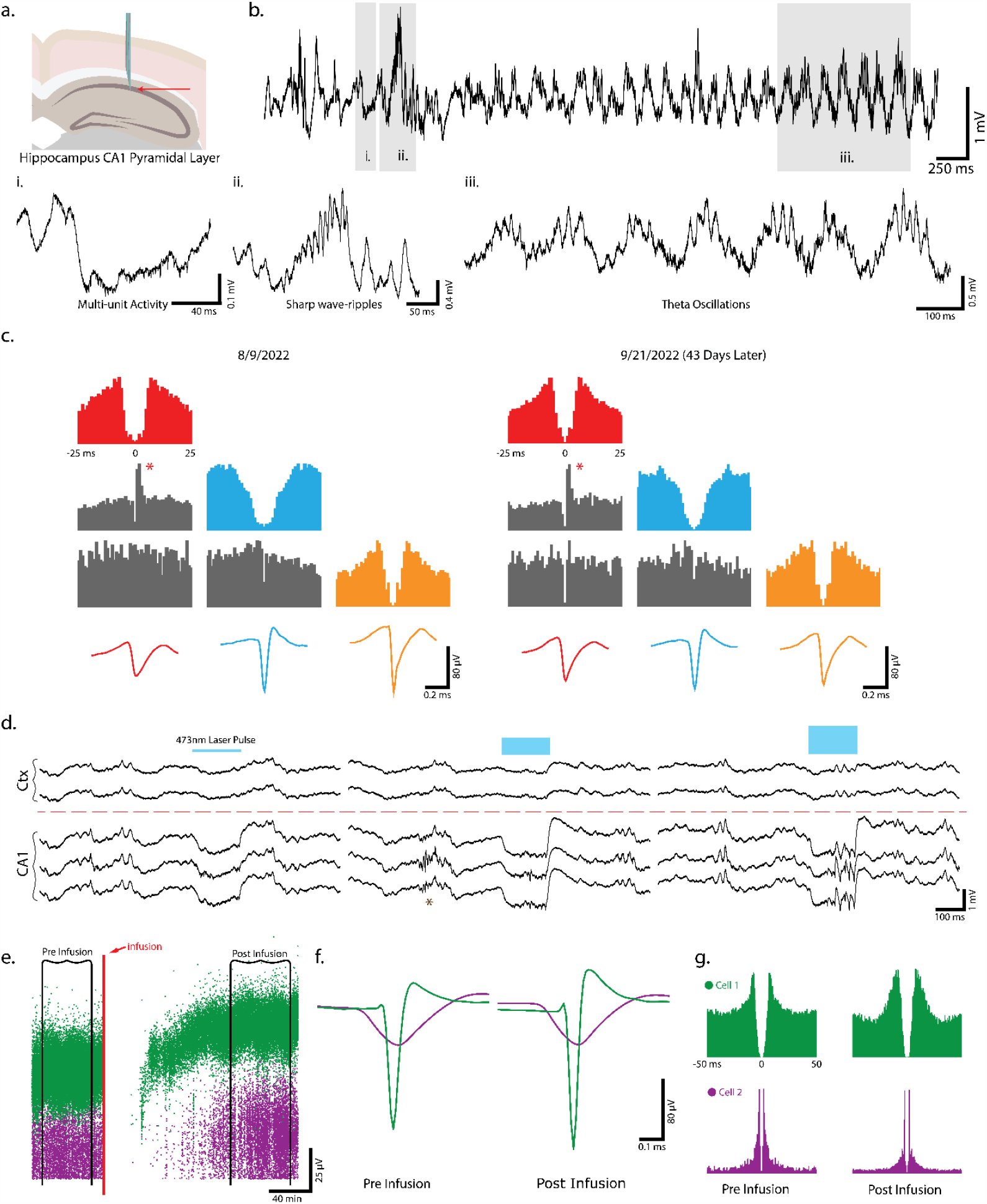
**a**. Schematic of the targeted implanted site, hippocampus CA1. **b**. Example wideband (0.1 – 8000 Hz) extracellular traces obtained from CA1. **(i)** Enlarged trace to display multi-unit activity. **(ii)** An example trace of a sharp wave-ripple (100-250 Hz) above the pyramidal layer. **(iii)** An example of theta-nested gamma oscillations (6-11 Hz; 40-80 Hz). **c**. Three example cells recorded on 8/9 and 9/21 (PI’s birthday). Each cell is color coded to match the auto-correlations and the spike waveforms. *Peak in cross-correlation across the red and blue cells suggest their excitatory monosynaptic connection is maintained over 43 days. **d**. Examples of optically evoked responses with three levels of laser power (units). No response was observed in the cortex due to AAV-ChR2 expression localized to CA1. The amount of optically evoked neuron activity can be increased by varying the light output, allowing us to achieve optically induced SPW-Rs at higher laser powers. * shows spontaneous SPW-Rs in between optical stimulations. **e**. Raster plot of an interneuron and a pyramidal cell to demonstrate the recovery after infusion (200nL; 1nL s^-1^). Given the device tip contains both the microfluidic channel and recording sites (< 20 μm in distance), the cells are pushed away and return after some period of time. **f**. Average spike waveforms of the two cells before and after infusion. **g**. Autocorrelation of the two cells before and after infusion.

We additionally chronically implanted the T-DOpE probes in CA1 (n=2 animals) to demonstrate biocompatibility and ability for long-term stable recording single unit recording capability. Three example putative neurons (i.e., single units) were identified from recordings on 8/9 and 9/21. Each cell is color coded to match the auto-correlations and the spike waveforms across days. The average waveforms recorded from each electrode are included in **Supplementary fig. 2**. The neurons can be classified by the shape of the waveform and autocorrelation^46^ (putative ID, red: pyramidal cell, blue: interneuron, and orange: pyramidal cell). The waveforms from each electrode and the auto- and cross-correlation suggest that the probe recorded from the same group of neurons across 43 days. Additionally, the peak in the cross-correlation from the red and blue cells (**fig. 4c\***) shows a monosynaptic connection where the red unit is driving the blue unit to fire action potentials^47, 48^. The observation that this monosynaptic connection maintained across 43 days further supports that fact that the probe can reliably record from the same population of neurons over at least one month, a common goal in modern probe designs^3, 4, 49^. These data demonstrate that our novel T-DOpE probe has sufficient long-term biocompatibility and stability to perform longitudinal experiments without disrupting local circuitry.

To validate the optical waveguides in our T-DOpE probe, we recorded from CA1 in mice with ChR2 expression restricted to CA1 pyramidal neurons. In these experiments, the probe is advanced until the four most ventral electrodes are ~100 um above the center of the CA1 pyramidal layer, with the four most dorsal electrodes remaining in overlying cortex. **Fig. 4d** presents examples of local field potential responses to three levels of optical stimulation intensity (i.e., laser power). No optically induced responses were observed in cortex, as expected because ChR2 expression was localized to CA1. In CA1, optical stimulation induced large negative deflections in the LFP, likely due to cation influx through ChR2 and active membrane conductances in the ChR2 expressing neurons, as well as synaptically evoked inward currents in postsynaptic neurons activated by glutamate release from the ChR2+ cells. By regulating optical stimulation power, we were able to modulate multi-unit spike rate (**fig. 4d**). Optogenetic activation of CA1 PYRs induced local high frequency ripple-frequency oscillations in CA1^50^. **Fig. 4d\*** indicates a SPW-R event endogenously occurring between optical stimulations. These findings demonstrate the probe’s ability to monitor and manipulate local circuits while preserving physiological functions of the local circuit.

There is a general problem in drug infusion into the brain in that tissue is displaced, which is especially noticeable when combined with single unit electrophysiology. Supporting the hypothesis that physical cell displacements, not pharmacological influence, induce this transient apparent silencing of cells, we observed transient absences of spikes after infusion of either saline, vehicle, or drug+vehicle (**Supplementary fig. 3**). **Fig**.**4e**, shows an example session where we infused 200 nL at 1nLs^-1^ infusion rate, and recording one putative pyramidal cell (purple) and interneuron (green). After the displaced tissue relaxed to its original position spikes were once again able to be detected, at the level of single both sorted cells (**fig. 4e**, post-infusion) and multi-unit activity (data not shown). **Fig. 4f,g** shows the spike waveforms and autocorrelations of the two cells before and after infusion. The similarities in their waveforms and auto-correlations suggest that these are the same interneuron and pyramidal cell. Importantly, small changes in waveform shape are likely due to the displacement and return of the tissue to the same location with some spatial jitter and thus a change in the effective resistance through the Ohmic brain tissue. However, this jitter is likely in the range of tens not hundreds of microns, explaining the similarity in baseline and recovery spikes. Over experiments the mean return time was 32±14 minutes (session number = 8). Of note, this is a major improvement over experimental designs which require drug infusion distant from recording sites, making the locus of effect difficult to interpret.

### CA1 theta power during running is reduced by pharmacological activation of CB1Rs expressed in CA1

To examine the role of CB1R in hippocampal CA1 activity during behavior, we used the T-DOpE probe to focally infuse CB1R agonist, CP-55,940, in CA1 while simultaneously monitoring neural activity (animal number = 6 awake head fixed mice). **Fig. 5a** displays an illustration of the experimental setup. Head-fixed mice ran on a vertical wheel driving a 1-D visual virtual reality environment that reliably promotes natural (i.e. spontaneous not due to training) running behavior. Fig. 5b shows a representative session of CP-55,940 infusion (16.8ng; 200nL; 1nLs^-1^) following one-hour recording of baseline (**fig. 5b,c,f** is from one CP-55,940 session). The velocity of the mouse is shown above the spectrogram of the session. The local infusion of CP-55,940 was sufficient to lower the power of theta oscillations in the LFP (calculated from data restricted to running epochs). The normalized power spectrum of baseline (Before infusion; duration: 60 minutes) and after drug infusion (one hour after infusion; duration: 60 minutes) are shown in **fig. 5c**. Note the peak at 8 Hz decreased by approximately 20%. **Fig. 5d** shows an example control session with drug vehicle (VEH), the normalized spectrum of the baseline and after VEH infusion. The comparison between baseline and VEH shows no significance while the comparison between the baseline and CP-55940 showed significant difference (Student’s two-tailed t-test, NS: not significant p≥0.05, * p≤0.05(Baseline vs. VEH, p=0.9117, animal number=3), (Baseline vs. CP-55,940, p=0.0134, animal number=3)) (**Fig. 5e**). Across all sessions (animal number=3 VEH, animal number=3 CP-55,940), theta power significantly decreased following CP-55,940 infusion compared to control. To the best of our knowledge, this is the first demonstration that focal CA1, as opposed to systemic^37^, agonism of CB1Rs is sufficient to reduce theta oscillations in CA1. These findings suggest cannabinoids can impair hippocampal oscillations by acting upon only CB1Rs expressed in CA1 (either Schaffer Collateral terminals from CA3, or local CA1 neurons), without acting on other brain areas.

**Figure 5:**
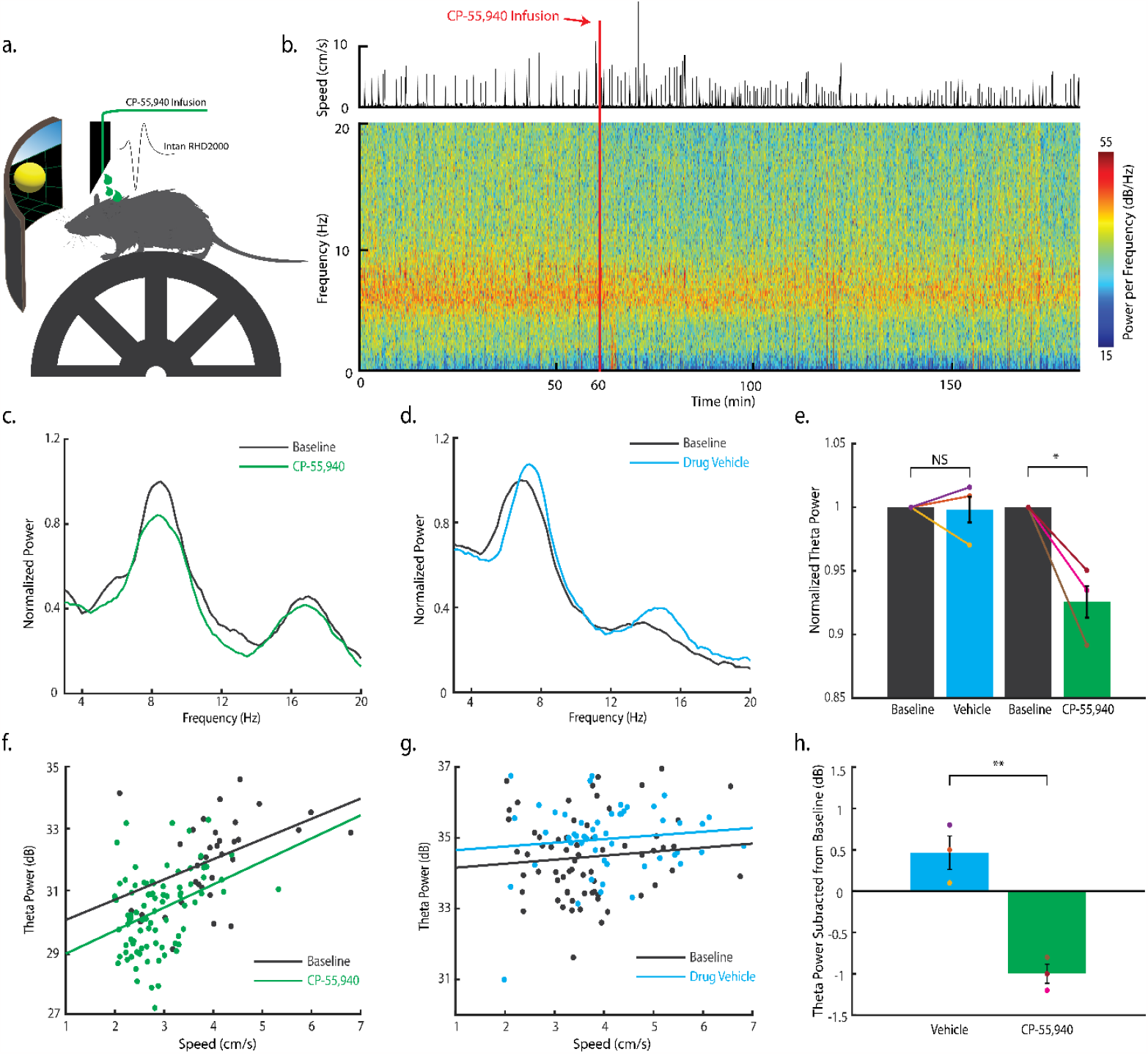
**a**. Illustration of the experimental setup. A head-fixed, wild-type mouse is mounted on the wheel. A virtual reality environment is presented on the screen to instigate running, and the virtual position is recorded. Local infusion and neural recordings are achieved using the T-DOpE probe. **b**. Power spectrogram of CP-55,940 infusion session in lower frequency bins (0 – 20 Hz). The upper panel shows the velocity of the mouse. CP-55,940 (mg kg^-1^; 200nL; 1nL s^-1^) is locally infused at the recording sites after 60 minutes of baseline. **c**. Normalized power spectrum of the baseline (Before infusion; duration: 60 minutes) and after CP-55,940 infusion (one hour after infusion; duration: 60 minutes) restricted to running time (>2cm s^-1^ and >1 s running epochs). **d**. Normalized power spectrum of the baseline (Before infusion; duration: 60 minutes) and after drug vehicle infusion (one hour after infusion; duration: 60 minutes) restricted to running time with the same criteria as above. **e**. Comparison of the normalized theta power (6 -11 Hz) between baseline and infused vehicle or drug (Student’s two-tailed t-test, NS: not significant p≥0.05, * p≤0.05(Baseline vs. Drug Vehicle, p=0.9117, animal number=3), (Baseline vs. CP-55,940, p=0.0134, animal number=3)). **f**. Correlation between theta power and the running speed during baseline and after CP-55,940 infusion. **g**. Correlation between theta power and the running speed during baseline and after drug vehicle infusion. **h**. theta power shift from the baseline to the vehicle or drug for all the sessions. (Student’s two-tailed t-test, ** p≤0.01(Vehicle vs. CP-55,940, p=0.0033, animal number =6))

In some sessions, the mouse ran slower after CP-55940 infusion. It is established that hippocampal theta power is positively correlated with running speed.^51-53^ It is possible that theta power reduction following drug infusion may in part be due to changes in behavior (e.g. CP-55,940 causes the mouse to run slower). Previous works with systemic CB1R agonist infusion have established that CB1R agonists reduce theta power beyond that explained by the decrease in running speed^37^. To investigate this with focal infusion, we calculated the correlation between the theta power and the running speed (**fig. 5f**) and found even when matched for running speed, theta power was lower after agonist infusion. **Fig. 5g** shows the correlation between theta power and running speed in the baseline and after VEH infusion. In **fig. 5h**, the theta power shifts were quantified for both VEH and CP-55,940 sessions (Student’s two-tailed t-test, ** p≤0.01(Vehicle vs. CP-55,940, p=0.0033, animal number=6)) The theta power increased from the baseline after VEH infusion; theta power decreased after CP-55940 infusion. Therefore, the local infusion of CP-55,940 is sufficient to reduce the power of theta band (6-11 Hz) compared to VEH, while running at the same velocity. This is a novel finding as systemic application of the drug affects subcortical and other cortical areas involved in theta generation, while here we demonstrate that CA1 CB1R agonism alone reduces CA1 theta.

### SWR rate is lowered by pharmacological activation of CB1Rs expressed in CA1

Systemic administration of CP-55,940 decreases SPW-R rate and power in behaving animals,^37^ and that under urethane anesthesia, intrahippocampal infusion of CP-55,940 reduces SPW-R power.^37^ Interestingly, we find similar results for local intrahippocampal infusion in CA1 of behaving mice, further implicating local CB1R expression in changes in CA1. With data recorded from the same setup shown in **fig. 5a**, sessions were analyzed in higher frequency bands (100-300 Hz) to study the influence of CP-55,940 on SPW-Rs. In **fig. 6a**, the power spectrogram of a CP-55,940 infusion session was computed with the ripple rate plotted on top. The upper panel shows the velocity of the mouse. CP-55,940 (16.8ng; 200nL; 1nLs^-1^) was locally infused at the recording site after 60 minutes of baseline recordings. After the infusion, the power in 100-300 Hz bins significantly decreased, which indicates a reduction in SPW-Rs. Importantly, ripple rate returned to baseline after ~ 6 hours. (**fig. 6b**). Spectral analysis of the baseline (Before infusion; duration: 60 minutes) after CP-55,940 infusion (one hour after infusion; duration: 60 minutes) were computed. The bump centered at 150 Hz in the baseline spectrum (**Fig. 6c**) disappeared following CP-55,940 infusion, due to the abolishment of SPW-Rs. The difference of the two spectra was plotted to further visualize the change in the power at each frequency (**fig. 6d**). The bump in **fig. 6c** is highlighted further by the negative peak at 150 Hz, due to impaired SPW-R generation. The ripple rate (per minute, normalized to ripple rate of the first hour) was calculated to investigate the impact of drug infusion on SPW-R occurrence (**fig. 6e**, animal number=7). The normalized ripple rate after CP-55,940 infusion was significantly reduced (Student’s two-tailed t-test, **** p≤0.0001(Baseline vs. CP-55,940, p=9.7829 e-48, animal number=3)) compared to VEH control. We compared the normalized SPW-R count between baseline and infused vehicle or drug in **fig. 6f**. (Student’s two-tailed t-test, **** p≤0.0001, * p≤0.05 (Baseline vs. VEH, p=0.0265, animal number=4), (Baseline vs. CP-55,940, p=9.7829e-48, animal number=3)). Overall, the injection of CP-55,940 (but not VEH) significantly lowered the ripple rate. This demonstrates for the first time the importance of local CA1 CB1R signaling in SPW-R dynamics in behaving mice.

**Figure 6:**
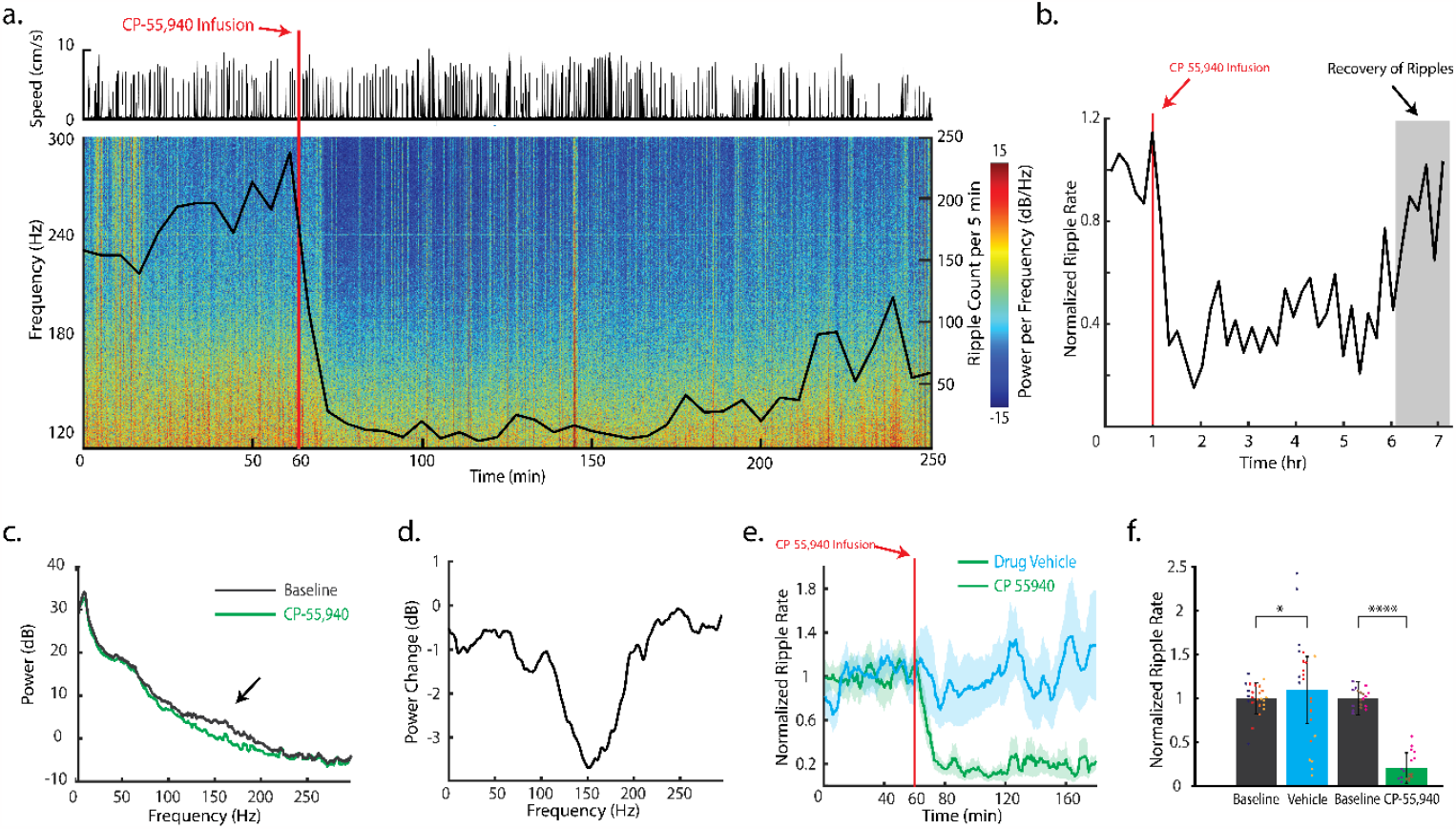
**a**. Power spectrogram of CP-55,940 infusion session on higher frequency bins (100 – 300 Hz) with the same experimental set-up as Fig. 5a. The ripple rate is plotted on top of the spectrogram with respect to the same timescale. The upper panel shows the velocity of the mouse. CP-55,940 (mg kg^-1^; 200nL; 1nL s^-1^) is locally infused at the recording sites after 60 minutes of baseline. **b**. Ripple count per 10 minutes over 7-hour recording session **c**. Power spectrum of the baseline (Before infusion; duration: 60 minutes) and after CP-55,940 infusion (one hour after infusion; duration: 60 minutes). **d**. Spectral difference from the baseline to the recording after CP-55,940 infusion. **e**. Normalized ripple rate across all animals and sessions (drug vehicle and CP,55-940 infusion; animal number = 7). **f**. Normalized ripple count between baseline and infused vehicle or drug (Student’s two-tailed t-test, **** p≤0.0001, * p≤0.05 (Baseline vs. vehicle, p=0.0265, animal number=4), (Baseline vs. CP-55,940, p=9.7829 e-48, animal number=3)).

### Generation of SPW-Rs by optical stimulation of CA1 PYR is abolished by pharmacological activation of CA1 CB1Rs

SPW-Rs are generated when CA1 receives strong excitatory input from CA3, with the sharp wave resulting from dendritic depolarization of CA1 PYR by CA3, and the fast oscillation due to intra-CA1 interactions between excitatory pyramidal neurons and local GABAergic inhibitory interneurons^30, 50, 54^. While our results confirm that spontaneous SPW-R rate decreases following intrahippocampal CP-55,940 infusion, it has remained unclear if this is due to signaling at CB1Rs expressed on synaptic terminals originating from CA3 inputs^43, 44, 55^ and/or by hippocampal interneurons. To address whether changes in CA3 inputs could explain the reduction in SWRs, we substituted the CA3 sharp wave excitation of CA1 PYR with direct ChR2 depolarization^50^. We used the T-Dope probe to optogenetically depolarize ChR2+ CA1 PYR before and after CP-55,940 infusion. Similar to the setup in **fig. 5a**, a head-fixed AAV-CamKII-ChR2 mouse was placed on the wheel with a 473nm laser connected to the T-DOpE probe to deliver optical pulses into CA1 (**fig. 7a**). Following 30 minutes of baseline, CA1 pyramidal cells were optically stimulated with a 150 ms optical pulses at low, medium, and high power over 40 minutes (**fig. 7b**; n=400 stimulations). 10 minutes after the last optical stimulation, CP-55,940 (16.8ng; 200nL; 1nLs^-1^) was focally infused. Once the tissue recovered from infusion, another set of optical pulses was delivered, using the same optical power levels. The spectrogram of the session was computed with the ripple rate overlayed, as shown in **fig. 7b**. Note the increased SPW-R events detected from 40–70 minute mark due to the optically induced SPW-Rs. After CP-55,940 infusion, the occurrence of SPW-R events drastically decreased and identical low, medium, and high light pulses were not sufficient to optically induce SPW-Rs (**fig. 7c**). Optical stimulation activated the network to the same extent before and after, as indicated by the fact that the mean optical LFP response to identical light pulses was not different (**fig. 7d**). Importantly, this demonstrates that failure to generate SPW-Rs post drug is not due to an inability to sufficiently depolarize CA1 neurons.

**Figure 7:**
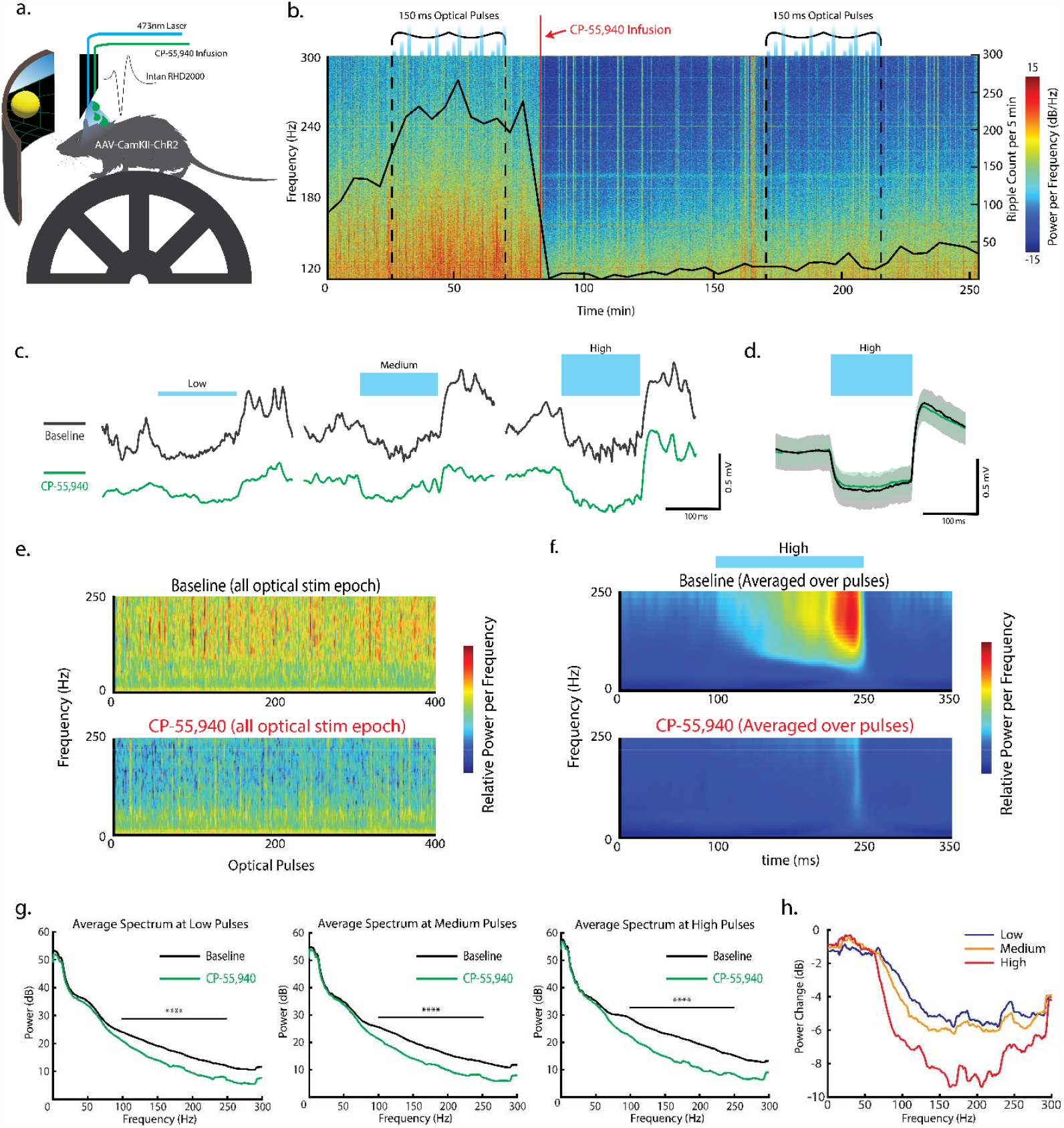
**a**. Illustration of the experimental setup. A head-fixed AAV-CamKII-ChR2 mouse is mounted on the wheel. A virtual reality environment is presented on the screen to instigate running, and the virtual position is recorded. Local infusion, optical stimulation, and neural recording are achieved using the T-DOpE probe. **b**. Power spectrogram of CP-55,940 infusion session on high frequency bins (100 – 300 Hz). Following 30 minutes of baseline, CA1 PYRs are optical stimulated (n=400 pulses of 150 ms at low, medium, and high power) over 40 minutes. CP-55,940 (mg kg^-1^; 200nL; 1nLs^-1^) is locally infused at the 80-minute mark. After the cells recover from the infusion, optical stimulation is repeated. **c**. Representative response to low, medium, and high optical stimulations during the baseline and after CP-55,940 infusion. **d**. Average neural response to the high optical stimulation during the baseline and after CP-55,940 infusion. **e**. Pseudocolor plots of the power spectra of high optical pulse evoked activities in the baseline and after drug infusion. **f**. Average Wavelet Transform of the activities in baseline and after drug invusion. **g**. Average spectrum of baseline and for CP-55,940 infusion during low, medium, and high optical pulses. **h**. Spectral difference between the baseline and after CP-55,940 infusion for the optical pulses.

To investigate the variance in power of induced SPW-R for individual optical pulses, we computed the spectrum for each pulse, displayed as pseudocolor plots (**fig. 7e** and **Supplementary fig. 4,5**). Wavelet transform was additionally computed and averaged over all sessions to visualized to optically induced SPW-Rs (**fig. 7f** and **Supplementary fig. 6,7**). For all optical pulses, significantly higher power was observed in the SPW-R band (100 – 250 Hz; (Wilcoxon signed rank test, **** p≤0.0001 (Baseline vs. CP-55,940, p<10^−32^ for all frequency bins in 100-250 Hz, n=400))) before drug infusion. **Fig. 7g** shows the average spectrum at low, medium and high optical pulses. The power in the 60-100 Hz range in the average spectra from the baseline progressively increases as the optical power increases. The differences between baseline and post-drug infusion were larger at higher optical pulse intensity (**fig. 7h**). These data further support agonism of CB1Rs expressed by CA1 interneurons as the mechanism of agonist induced SPW-R suppression.

## Discussion

Here we fabricated novel T-DOpE probes via a scalable and low-cost thermal tapering method. The method enables high versatility and complexity in probe design, and the semi-automated connection method ensures scalability of the T-DOpE probes. Our *in vivo* study demonstrates promising future experiments with the T-DOpE probe’s reliable and precise recordings with simultaneous optical, and chemical manipulations to understand complex neural circuitry. Here, we used T-DOpE probes to investigate the role of CA1 CB1R signaling in hippocampal activity including theta and SPW-Rs in behaving animals. Our results suggest a critical role for CB1R signaling in the ability for CA1 circuits to generate ripples.

Typically, it is desirable to achieve a thinner probe to minimize tissue response^56^ (i.e., neuron-scale devices). The thermal tapering process accomplishes this goal by allowing us to make an ultrafine tip at the tissue interface while maintaining a sizable backend that is compatible with industrial-level soldering/bonding process. For example, we can easily draw down mini-preforms to produce a 50 μm sensing tip with a backend diameter of 2 mm. Compared with cleanroom microfabrication-enabled devices, our approach allows low cost fabrication of flexible and multifunctional devices. Compared with fiber probes fabricated using TDPs, our T-DOpE devices enable dual-size end tips instead of a consistent diameter. In previous fiber probes, electrodes were manually connected by carefully scraping away insulated layers and then electrically connecting the exposed electrode to a pin. Microfluidic channels are connected by attaching polymer tubing to the fiber via a similar process as the electrode connection. Optical connection is done by attaching a ferrule to the back of the fiber, limiting it to one optical connection per device. This manual connection is a time-consuming and labor-intensive process that becomes increasingly difficult as the fiber gets thinner. In our T-DOpE probes, the relatively large size of the backend allows for a semi-automated connection process, reducing connection time, labor, and cost (**fig. 3a**; Cost: <10 dollar per unit). In addition, the previous connection process only allows connections to the outer layers of the device, and it is impossible to connect the channels near the center of the fiber without damaging outer channels. This restricts the complexity of the previous probes. Our new connection process enables electrical, optical, and chemical modalities in the entirety of the fiber to be easily connectorized and also allows T-DOpE probes to be readily available for wide distribution.

Using our newly developed device, we demonstrated precise optical and chemical modulations *in vivo*. We successfully identified the same monosynaptic connection between a putative pyramidal cell and an interneuron across 43 days which highlights the stability of long-term recording (**fig. 4c**). The biocompatibility of the materials in T-DOpE probes (i.e., BiSn, PC, and PVDF) have been previously established.^23, 26^ By carefully varying the optical power, we were able to achieve different levels of manipulations of neural circuitry: increasing firing rate and optically inducing SPW-Rs (**fig. 7c, 7e**). We also characterized the push and return of neurons after local infusion via electrophysiology (**fig. 4e**). The slow local infusion allows us to study the pharmaceutical/chemical effects of a local neural circuitry. The slanted cut of the tapered probe minimizes the tissue damage during implantation and enables depth-dependent recording along the device tip. With the T-DOpE probes, we always recorded the typical electrophysiological landmarks of hippocampus CA1, such as multi-units, SPW-Rs, and theta oscillations (**fig. 4b**). The performance and reliability of our T-DOpE probes is comparable and, in certain aspects, superior to commercially available silicon neural probes.

The hippocampus plays a major role in spatial memory. During exploration, CA1 pyramidal place cell (PYR) activity is organized at the theta timescale (6-11 Hz)^33, 34, 57^. During consummatory behaviors and non-rapid eye movement sleep, large depolarization events drive CA1 PYR and local inhibitory interneurons to interact and lead to the generation of sharp wave-ripples (SPW-R, 100-250 Hz)^30-32^. Previous studies found that cannabinoids alter CA1 rhythms, suggesting a possible role for local CB1R in hippocampal activity and oscillations supporting memory^37, 42^. However, CB1Rs are expressed in multiple hippocampal sub-regions (e.g., both CA1 and CA3) and cell types (e.g., excitatory pyramidal cells vs inhibitory interneurons) ^43-45, 58, 59^, and the specific loci of action of cannabinoids on CA1 activity remains unknown.

It has been shown that administration of CB1R agonists is sufficient to disrupt theta while keeping neuronal firing rate intact. In a related study, the same group went on to show that temporal coding in the hippocampus is disrupted by cannabinoids^37, 42^. While it is clear that cannabinoids disrupt hippocampal circuitry coordination underlying theta, these studies were achieved through systemic administration of cannabinoids in rats. Until our study, it was unknown if cannabinoids were disrupting theta generated upstream of CA1, or within local circuitry. The medial septum, a proposed theta generator, has glutamatergic, cholinergic, and GABAergic connections to CA1^59^. It has been shown that GABAergic and cholinergic neurons express CB1 receptors, likely inhibiting vesicle release at axon terminals.^58, 60^ One hypothesis is that systemic administration of cannabinoids may decrease theta by disrupting circuitry within the medial septum, which prevents effective theta drive in CA1. An alternative hypothesis is that cannabinoids bind to CB1 receptors expressed in CA1, either CA1 localized neurons or terminals projecting into CA1, and alter precise timing of neurotransmitter release disrupting oscillations^42^. We demonstrated with our T-DOpE probe that local infusion of CP-55,940 was sufficient to disrupt theta (**fig. 5e**), while maintaining the relationship between velocity and theta power (**fig. 5h**). This is the first study in a behaving animal, that shows local CA1 cannabinoid infusion is sufficient to diminish theta power. This finding supports other studies, implicating local CB1R activation^35, 37^ in the disruption of spatial memory associated with cannabinoids, either in CA1 neurons, or terminals from areas such as the medial septum, as opposed to binding in areas where the oscillation is generated from.

SPW-Rs are an important electrophysiological marker of learning and memory. Elongating SPW-Rs leads to increased memory performance, while disrupting SPW-Rs diminishes memory.^30-32^ It has been demonstrated that cannabinoids inhibit SPW-Rs in rats^37^, offering a potential mechanism underlying the memory impairment associated with cannabinoid use in humans. In previous studies, hippocampal administration of CB1 agonists inhibit SPW-Rs, thought to be caused by reduced glutamate release in CA1. ^37, 61^ CB1Rs are expressed in many locations within CA1, making it unclear where cannabinoids are acting to inhibit SPW-R generation. Both CA3 and CA1 neurons express CB1Rs.^43-45, 58, 59^ CB1Rs on CA3 axon terminals of the Schaffer collaterals have been shown to be involved in decreased glutamate release.^43, 44^ CA1 PYR express CB1Rs localized on their dendrites, activation has been shown to decrease post-synaptic excitability by increasing membrane conductances.^62^ CB1R are highly enriched in CA1 INT, suggested to inhibit GABA release.^45, 58^ Given the ambiguous relationship of CB1R expression and SPW-R generation, it was unclear if cannabinoids were acting on excitatory drive from CA3 or local circuit interactions within CA1. Our novel device allows us to directly interrogate CA1 through combining localized drug delivery and optogenetic stimulation. Optogenetic activation of pyramidal cells expressing ChR2 in CA1 results in sufficient excitation to induce SPW-Rs (**fig. 7e**).^50^ The fact that the CA1 circuit cannot generate SPW-Rs under a CB1R agonist, even if the network is sufficiently activated (fig. 7d), suggests that the lack of ripple oscillations is due to activation of CB1R expressed on INT terminals: activation decreases the precision of GABA release and inhibition pacing PYR populations as is necessary for SPW-R generation. This strongly suggests that the agonist mediated disorganization of inhibitory inputs to pyramidal neurons is a primary mechanism by which cannabinoids suppress SPW-Rs.

## Method

### Preform Fabrication

For this study, we fabricated preforms with three different designs corresponding to the 3 different devices used. Preform fabrication for the thermal drawing process (TDP) is an iteration of four steps: machining, inlaying, film wrapping, and consolidation. For all the preforms in this paper, consolidation is performed by heating the preform under vacuum at 190°C. All polymer materials (films, tubes, rods, preforms and mini preforms) used during fabrication were baked under vacuum at 80°C to ensure they are moisture free. For the 8 electrode, 1 microfluidic channel and 1 waveguide fiber preform, we start by rolling PVDF films (McMaster-Carr) onto a PC tube (McMaster-Carr). We then rolled PC films (Laminated Plastics) onto the preform and consolidated it. 8 grooves were machined into the consolidated preform and inlayed with BiSn alloy (Indium Corporation). Additional PC films were wrapped around the preform, which is then consolidated. The 8 electrode, 8 microfluidic channel and 4 waveguide fiber preform was fabricated by first milling 4 hollow channels into a solid PC rod. The preform was then wrapped with PC film and consolidated. 4 more hollow channels were machined, and the preform was again wrapped with PC film and consolidated. 12 more hollow channels were machined into the resulting preform, and were inlayed with 8 BiSn alloy strips (Indium Corporation) and 4 polymer waveguides (PC core, PMMA cladding). Finally, the preform was wrapped with additional PC film and consolidated. The 12 microfluidic channel, 8 waveguide fiber was fabricated in a very similar manner to the process described above. The only differences being 8 hollow channels were machined for the second layer of microfluidic channels, and there were 8 waveguides instead of 4 and no BiSn strips inlayed in the outermost layer of device feature elements.

### Mini-preform Fabrication

Thermal drawing was accomplished by heating the finished preform in a custom-built furnace and pulling it down into a fiber using a capstan motor. The furnace is divided into top, middle and bottom sections that can be individually set to different temperatures. The top section preheats the preform, the middle section softens the preform and is where the fiber is drawn. The bottom section is where the fiber is cooled. For our fabrication process, the furnace’s sections were set at 150°C, 275°C and 120°C respectively (Note these may not be the actual preform temperatures as they are readouts of the temperature sensors mounted in the furnace). The drawn down fiber’s diameter is maintained at 2mm. The fiber is cut into 10cm segments to form mini-preforms for thermal tapering process.

### Tapered Device Fabrication

Tapering was done using a custom-built thermal tapering setup. The mini-preform was held in place and the alignment adjusted using Thorlabs opto-mechanical components. A DC power supply was then used to power a custom-designed furnace to 230°C. Note this is the nominal temperature measured by a thermal coupler and may not be the mini-preform’s actual temperature. Once the mini-preform was softened, it was pulled via a computer controlled linear motor (Zaber Technologies), resulting in a tapered structure. The speed of the motor and its travel distance can be utilized to adjust the tapered structure’s geometry. To produce a neural probe which minimally damages the nearby tissue, the tapered structure was cut at an angle, resulting in two individual T-DOpE probes.

### Assembly of Multifunctional tapered microprobe

The connection of the T-DOpE probe was accomplished on a custom connection setup built with opto-mechanical components. Translation stages (Thorlabs) and a digital microscope (Linkmicro) were used to provide finer position control during connection. For electrical connection, the device backend was heated to 160°C. This temperature is high enough to melt the BiSn electrodes but low enough to not damage the probe. 42AWG copper wires (Remington Industries) guided by a hypodermic needle were lowered into the melted electrodes. The device was then cooled to solidify the BiSN electrodes. Microfluidic connection was accomplished by inserting a custom drawn hollow (150 μm OD, 75 μm ID) PC tube into the microfluidic channels on the probe’s back end. Thermally drawn 200 μm diameter Polymer optical waveguides (PC core, PMMA cladding) were polished (30-1 μm grit) and carefully coupled onto waveguides on the probe’s connection end. To seal the microfluidic connection and keep the waveguide in place, UV resin (Piccassio) was applied to the entire probe backend and cured.

To properly interface with our recording setup, optical laser, and drug delivery systems, the probe is further fitted with adapters components. The probe’s copper wires were soldered to either pin connectors (chronic implantation) or custom designed PCBs (acute implantation) which can be readily connected to Intan acquisition systems via PCBs purchased from NeuroNexus. The probe’s waveguides were connected to Ø1.25mm stainless steel ferrules (Thorlabs). The microfluidic tubes were connected to IDEX Health & Science fluidic components via UV resin. The IDEX components themselves were made compatible with our drug delivery system by a custom-made adapter.

### Electrochemical spectral impedance measurement

Impedance measurements are done via a potentiostat (Gamry Instruments). The measurements were performed by lowering the sensing end of a fully connected T-DOpE probe into phosphate-buffered saline (PBS, Thermo Fisher). A Pt wire (Basi) was used as a counter and reference electrode. Data acquisition was accomplished via Gamry’s proprietary software.

### Headbar Implantation

Mice are induced and maintained at a surgical plane of anesthesia with isoflurane while mounted in stereotax. Hair is removed and scalp is disinfected. Bupivicaine nerve block is injected once under the scalp. The scalp is removed with surgical scissors and the skull is cleaned and dried with 3% hydrogen peroxide, followed by application of the sterile dental adhesive Optibond (Kerr Dental), which is cured with blue light. A < 0.2 mm burr hole is made above the right cerebellum, and a stainless-steel wire is inserted between the skull and the brain, parallel to the brain surface, then the wire is affixed to the skull with sterile dental acrylic. This wire is connected to the ground of the system. A titanium headplate (2 cm long, ~ 1 gram) is positioned above lambda, parallel to skull, and permanently fixed in place with sterile dental acrylic.

### Craniotomy

Mice are induced and maintained at a surgical plane of anesthesia with isoflurane while mounted in the stereotax. A 0.5-1.0 mm burr hole (using dental drill with 0.2 mm burr bit) is made above the hippocampus and the dura is removed. Biocompatible silicon elastomer (Kwik-cast; World Precision Instruments) is applied to the burr hole.

### AAV Injection

Mice are induced and maintained at a surgical plane of anesthesia with isoflurane while mounted in stereotax. Hair is removed and scalp is disinfected. Bupivicaine nerve block is injected once under the scalp. A < 0.2 mm burr hole is made above the hippocampus (mm from bregma: −1.8, lateral: 1.5). Glass pipette containing AAV5-CaMKIIa-hChR2(H134R)-EYFP (UNC Gene Therapy Center – Vector Core) is lowered into CA1 (mm from surface: -1.2). 100nL of AAV (titer: 4.1x10^12^ GC/mL) is injected into tissue (1nLs^-1^) using microinjector syringe pump (WPI: MICRO2T & 504127). Glass pipette is left for 5 minutes for virus to diffuse, then removed from brain. Craniotomy is covered using biocompatible silicon elastomer (Kwik-SIL; World Precision Instruments) and scalp is closed with Vetbond (3M).

### Drug

CP-55,940 stock was prepared in ethanol at a concentration of 1.68mg/mL. CP-55,940 injection solution was a mixture of 1:1:18 ethanol, solubilizer, and saline for a final drug concentration of 84μg/mL. 200nL of solution was delivered at a rate of 1nLs^-1^ directly into CA1. For a total delivery of an estimated 16.8ng or 44.6picomols. Drug vehicle solution was a mixture of 1:1:18 ethanol, solubilizer, and saline. 200nL of vehicle solution was delivered at a rate of 1nL s^-1^ directly into CA1. We used a standard precision injection apparatus (NanoFil Syringe and UMP-3 Syringe pump, World Precision Instruments) for all tests and experiment.

### *In Vivo* recording

Mice, trained and habituated to head-fixed navigation, are placed in the head fixation apparatus, and the T-DOpE probe is lowered through the craniotomy into the brain until SPW-Rs are recognized and left in place for 30-45 minutes until tissue is relaxed. The amplified neural signals are then recorded with RHD2000 system (Intan Technologies LLC). For chronic recordings, probe was fixed to the skull with dental cement, after the probe was lowered into CA1 and SPW-Rs were identified.

### Optical Stimulation

The T-DOpE probe’s optical ferrule was coupled to a diode-pumped solid-state (DPSS) laser (Laserglow Tehcnologies, 100mW maximum power, wavelength = 473nm.) through a mating sleeve (Thorlabs). We calibrated the optical output of our probe for each optical session. The optically evoked activity was closely monitored to meet the desired power output. The outputs of the optical power varied between 50 -700 μW. For investigating the effect of CP-55,940, 400, 150 ms optical pulses at low, medium, and high power were delivered. Time in between the power levels were set to 1 second (one set), and time in between stimulation sets was 5 seconds long.

### Data analysis

Data analysis was carried out with Matlab (The Mathworks) and custom scripts were used to analyze the extracellular recording. For each session, the LFP power analysis was performed on the electrode right above the electrode detecting highest power of SPW-R. The data was low-pass filtered and then downsampled from 30 kHz to 1250 Hz. Spectrograms were computed to visually analyze the sessions in both time and frequency domain. The calculations of spectrograms were computed using hamming window of 5 seconds and overlaps of 2.5 seconds. The calculation of power spectral density was performed by using the multi-taper estimate method. SPW-Rs were detected using the LFP data. The LFP was first spectrally filtered from 100 to 250 Hz and threshold filtered to 1.5-3.0 of the standard deviation of the whole data. The events that exceed 40-50 ms were counted towards the SPW-R events. The SPW-R was visually inspected and compared with the raw recording. The threshold and the duration of the SPW-Rs were manually adjusted to minimize the error in detection. The running epochs were computed by taking the derivative of the virtual position of the mouse (>2cm/s and >1s running epochs). The power spectra of the running epoch were also computed using the multi-taper estimate method. The spectra were also minmax normalized for the comparison over all the session (**fig. 5c,d**). Since the peak of the theta power varied over the animals and sessions, we computed the area under the curve (6-11 Hz) for the comparison between baseline and vehicle or drug (fig. 4e). The average running speed were computed for each running epoch. We used a linear fit for computing the correlation between the running speed and the theta power (**fig. 4f,g**). The difference between the linear fit between baseline and the drug/vehicle was computed for statistical comparison (**fig. 4h**). Spectra of the optically induced activity were computed using the multi-taper estimate. A pseudocolor plot was computed to visually represent the spectral analysis of all the pulses at once. To maintain the temporal information, the Wavelet Transform was computed and averaged. The LFP data was whitened for the pseudoplot and Wavelet Transform. Spike sorting was performed using Kilosort1, followed by manual refinement using Phy^63^. The cross-correlogram and auto-correlogram were normalized for visual observations. Experimental designs with two comparison groups were analyzed by two-tailed t-test with an assumption that the data is normally distributed. Wilcoxon signed rank test was used for the comparison of optically induced SPW-Rs in the baseline and after CP-55,940 infusion. The difference between the two groups were considered statistically significant if the p-value is less than 0.05. All tests were performed using Matlab code.

## Data availability

The raw data that support the findings of this study are available from the corresponding author upon request.

## Code availability

The MATLAB scripts for analysis are available from the corresponding author upon request.

## Acknowledgement

We thank Matthew Buczynski and Cristina Milano for advising us with the drug delivery of cannabinoids *in vivo*. X.J. gratefully acknowledges funding support from the National Institute of Health (R01NS123069, R21EY033080) and National Science Foundation (ECCS-1847436). DFE gratefully acknowledges funding support from The Simons Foundation and The Whitehall Foundation.

## Contributions

J.K., H.H., E.G., D.E., and X.J. designed the study. H.H. developed the tapering process. J.K. and H.H semi-automated the device connections. H.H. fabricated the T-DOpE probes used in the *in vivo* experiments. J.K. and E.G. carried out the animal surgeries. J.K. and H.H. optimized the electrophysiological recording, optical stimulation, and drug delivery setup. J.K. executed drug infusion experiments. J.K., H.H., and E.G. conducted simultaneous optical stimulation and drug infusion experiment. J.K., H.H., E.G., and K.A. analyzed electrophysiology data. All the authors contributed to the writing of the manuscript.

## Notes

### Competing Interest Statement

The authors have declared no competing interest.

