## Supplementary information for "Tapered Drug delivery, Optical stimulation, and Electrophysiology (T-DOpE) probes reveal the importance of cannabinoid signaling in hippocampal CA1 oscillations in behaving mice"

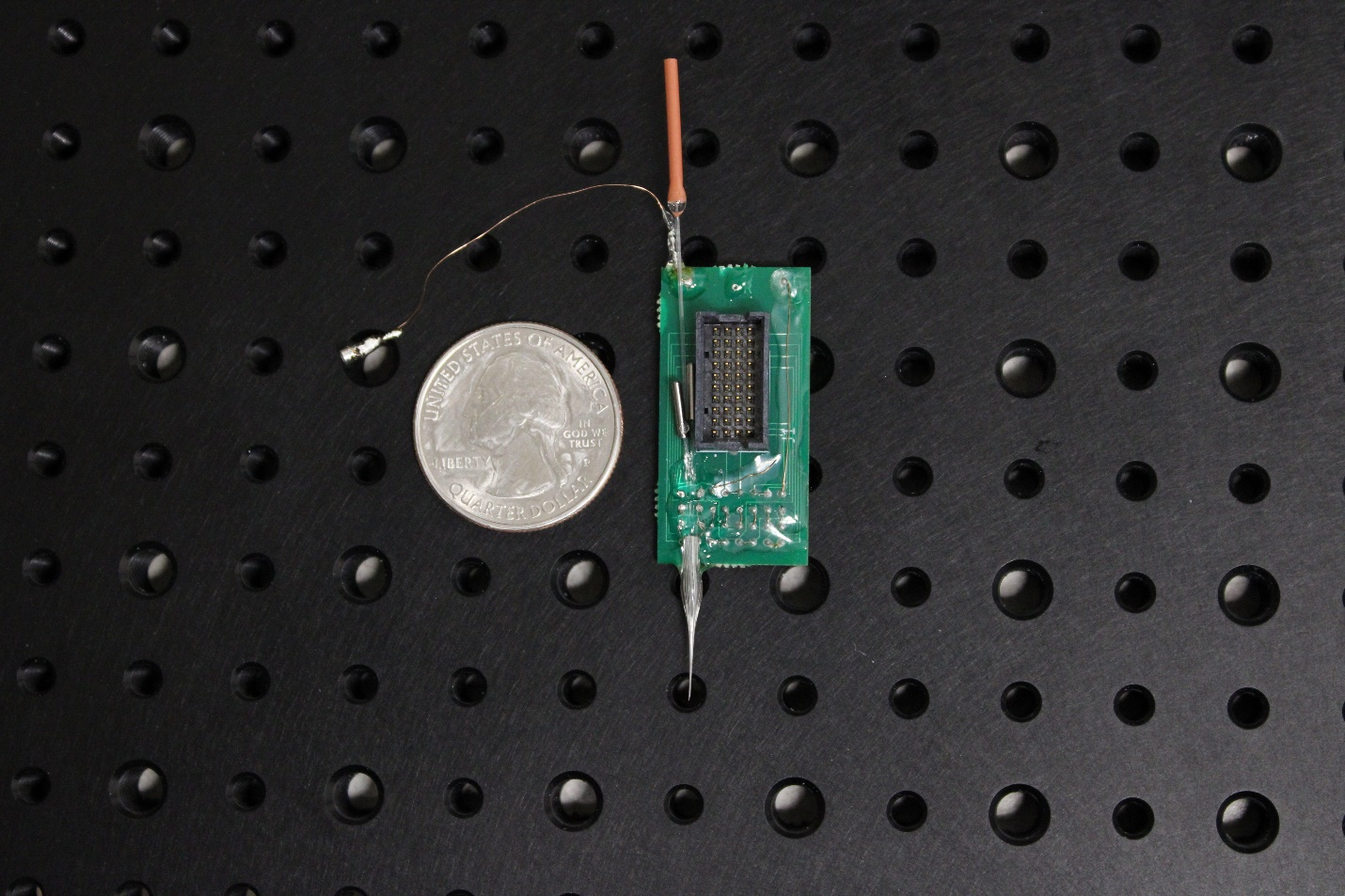

Supplementary Figure 1: Photograph of T-DOpE probe connected to PCB, ferrules, and microfluidic tube. T-DOpE probe connected to PCB was mainly used to monitor and manipulate CA1 circuitry acutely in behaving mice.

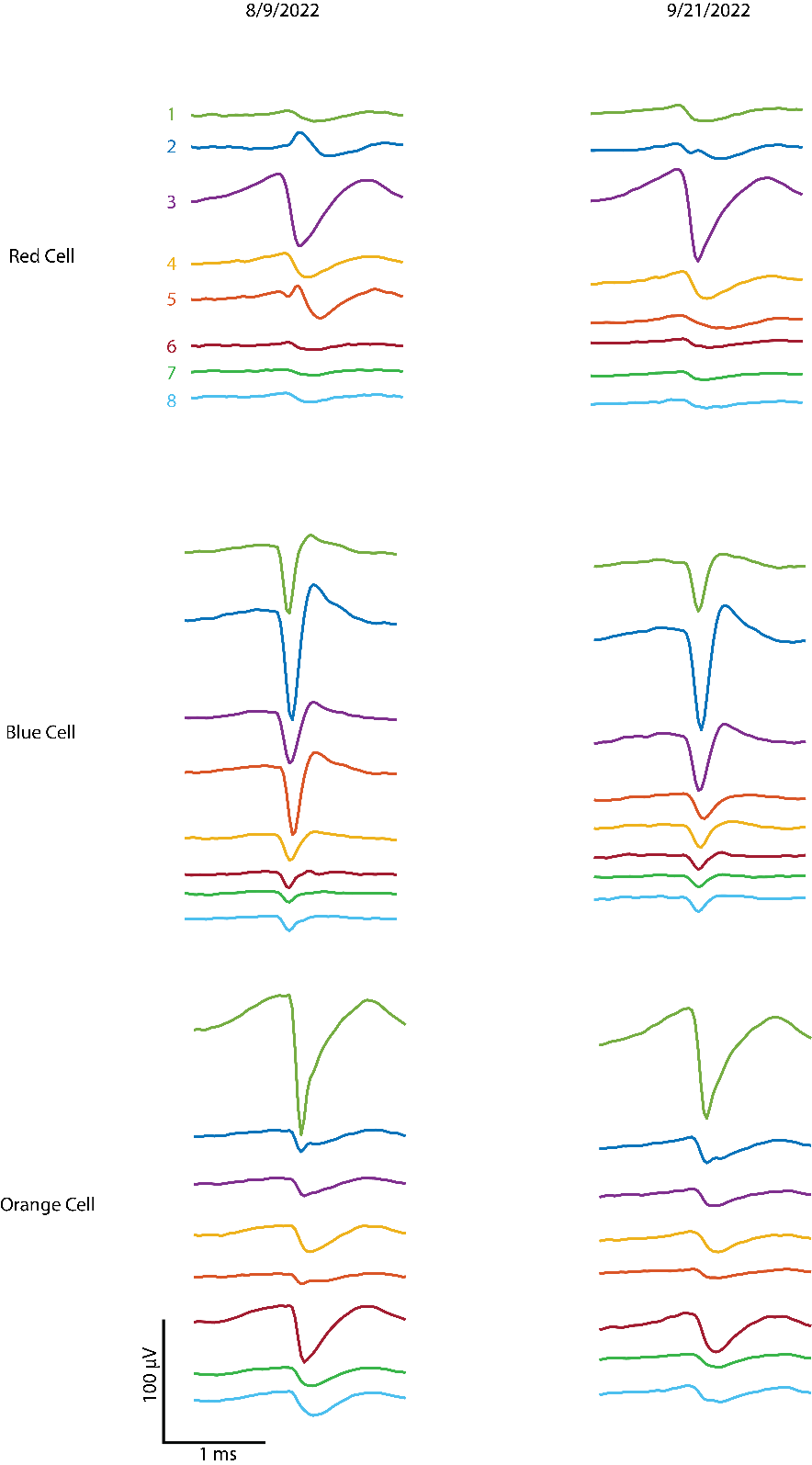

Supplementary Figure 2: Average waveform of identified cells in fig. 4c from each electrode. Red and blue cell maintained its monosynaptic connection over 43 days. The electrode number is color-coded and is not correlated with the location of the recording site.

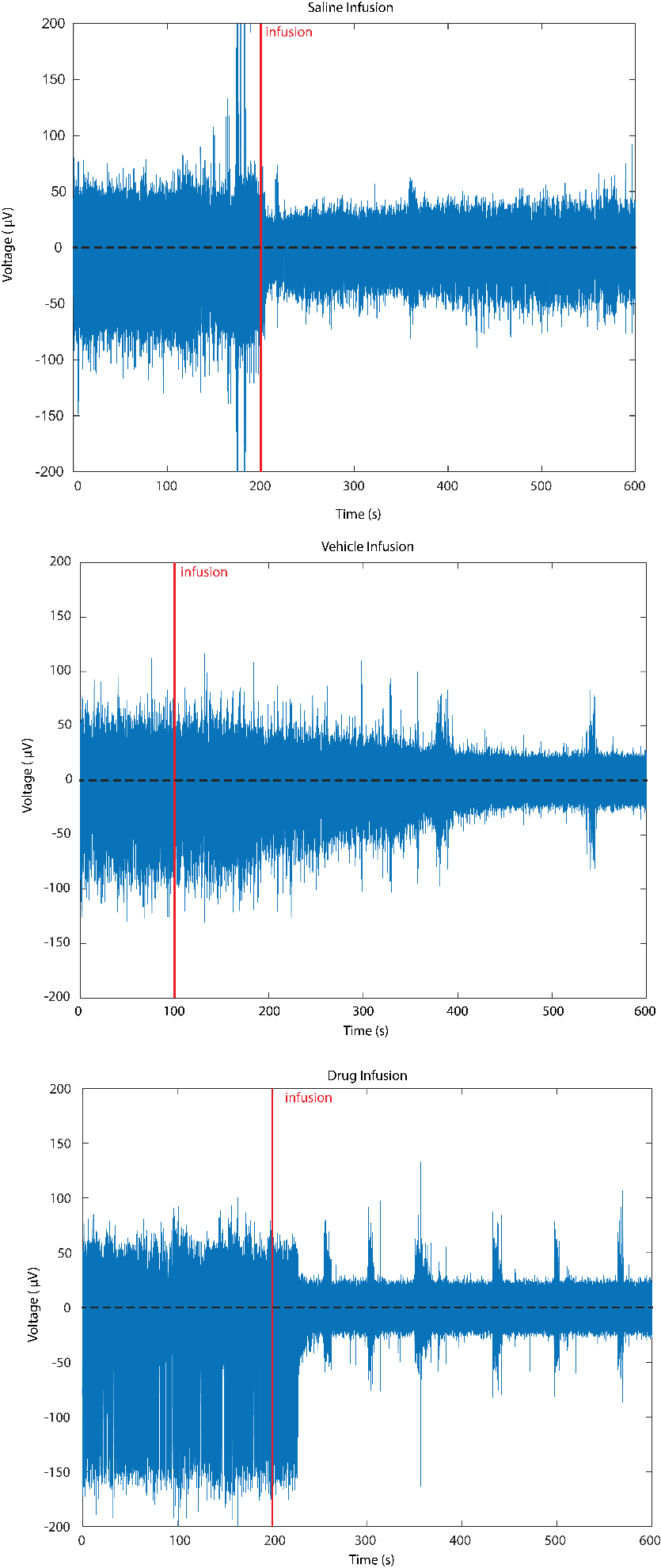

Supplementary Figure 3: 10-minute single-unit resolution extracellular trace (600- 8000Hz) where the cells are inevitably displaced due to infusion (200nL, 1nLs^-1^). Before infusion, the multi-unit activities were recorded with T-DOpE probe. For all infusion (saline, vehicle, and drug), the neurons are pushed away, and the action potentials of these neurons cannot be recorded until the diffusion has finished. The symmetric noise are the muscle artifacts from the mice.

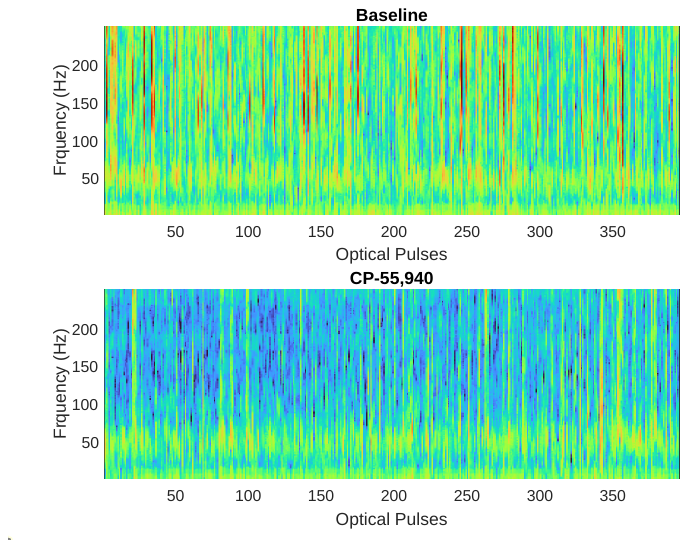

Supplementary Figure 4: Pseudocolor plot stacked with power spectral density of neural activity form each medium power light pulse (Top: before drug infusion, Bottom: after drug infusion).

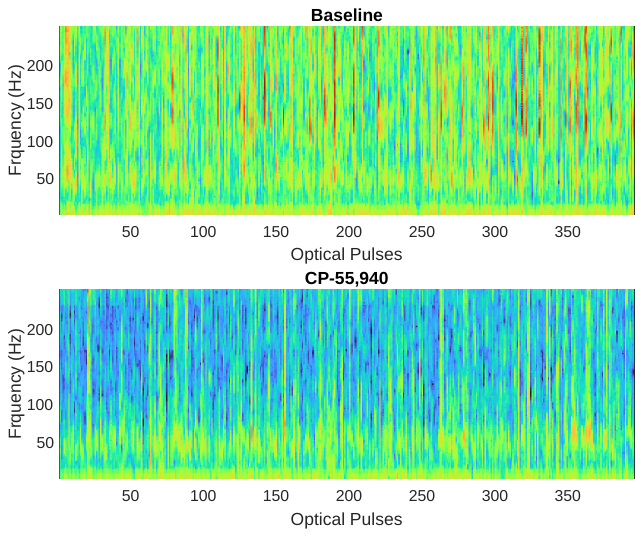

Supplementary Figure 5: Pseudocolor plot stacked with power spectral density of neural activity form each low power light pulse (Top: before drug infusion, Bottom: after drug infusion).

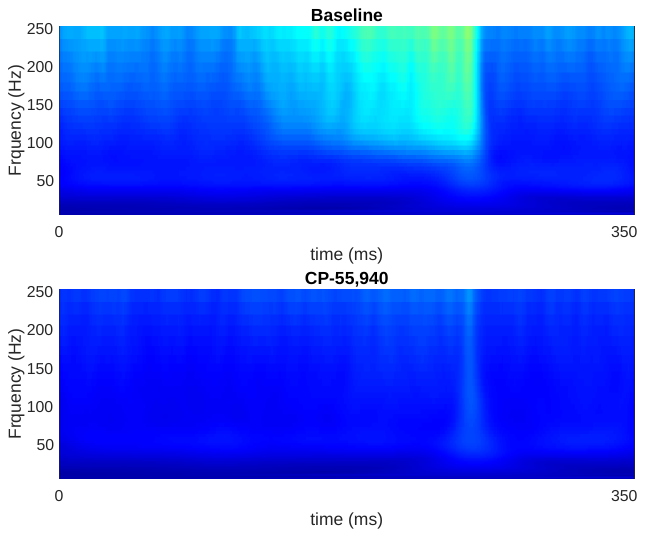

Supplementary Figure 6: Average of the Wavelet Transforms of LFP recording from 100 ms before optical stimulation to 100 ms after optical stimulation (150ms medium power optical stimulation; Top: before drug infusion, Bottom: after drug infusion).

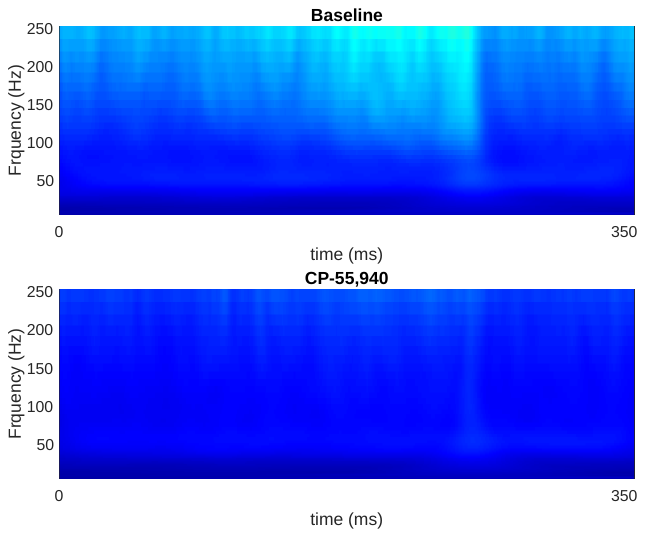

Supplementary Figure 7: Average of the Wavelet Transforms of LFP recording from 100 ms before optical stimulation to 100 ms after optical stimulation (150ms low power optical stimulation; Top: before drug infusion, Bottom: after drug infusion).
